# Chemogenetic dissection of the primate prefronto-subcortical pathways for working memory and decision-making

**DOI:** 10.1101/2021.02.01.429248

**Authors:** Kei Oyama, Yukiko Hori, Yuji Nagai, Naohisa Miyakawa, Koki Mimura, Toshiyuki Hirabayashi, Ken-ichi Inoue, Tetsuya Suhara, Masahiko Takada, Makoto Higuchi, Takafumi Minamimoto

## Abstract

The primate prefrontal cortex (PFC) is situated at the core of higher brain functions by linking and cooperating with the caudate nucleus (CD) and mediodorsal thalamus (MD) via neural circuits. However, the distinctive roles of these prefronto-subcortical pathways remain elusive. Combining in vivo neuronal projection mapping with chemogenetic synaptic silencing, we reversibly dissected key pathways from PFC to the CD and MD individually in single monkeys. We found that silencing the bilateral PFC-MD projections, but not the PFC-CD projections, impaired performance in a spatial working memory task. Conversely, silencing the unilateral PFC-CD projection, but not the PFC-MD projection, altered preference in a free-choice task. These results revealed dissociable roles of the prefronto-subcortical pathways in working memory and decision-making, representing the technical advantage of imaging-guided pathway-selective chemogenetic manipulation for dissecting neural circuits underlying cognitive functions in primates.

## Introduction

The prefrontal cortex (PFC) is well known to serve as the center of higher-order executive functions, making primates what they should be in terms of evolution (1). These functions, however, do not solely rely on PFC neurons, but on their cooperative actions with subcortical structures, including the caudate nucleus (CD) and mediodorsal thalamus (MD) (2–4). For example, working memory and decision-making — two fundamental functions that rely on the dorsolateral part of PFC — are impaired by lesions in either CD or MD in humans and non-human primates (5–8). Deficits in such cognitive functions have also been implicated in aberrant functional connectivity between PFC and CD/MD in patients with neurological, neuropsychiatric or substance abuse disorders (9–11). Despite growing evidence for the importance of PFC-derived cognitive signals in CD/MD, their causal contributions to working memory and decision-making remain unidentified in primates.

Anatomical studies have shown that the majority of the prefronto-subcortical projections arise from deep layers in monkeys (12, 13). In addition, PFC, MD and/or CD constitute two loop circuits, namely the cortico-basal ganglia-thalamo-cortical loop and the cortico-thalamo-cortical loop, which partially overlap (14). Accordingly, this raises the critical open question of whether the PFC to MD/CD pathways are involved in cognitive functions similarly or differently. To answer this question, it is essential to manipulate the prefronto-subcortical circuits independently.

Several studies have so far shown behavioral modification with reversible pathway-selective dissection in monkeys, where a double-viral vector infection method was used to manipulate the activity of a single neuronal pathway (15–18). This approach, however, is not suitable for investigating multiple neural pathways independently within a single animal. Optogenetics is another approach to manipulating the neuronal activity of a specific pathway by optical stimulation of axon-terminal sites. Although a few studies have succeeded in pathway-specific optogenetic activation in non-human primates (19–21), optogenetic silencing of synaptic transmission remains challenging (22, 23). Moreover, these approaches require precise identification of the locations of anatomically-connected multiple regions, i.e., viral injection or allocation of optic fiber, which is technically demanding when using non-human primates because of limited resources.

To elucidate the causal roles of distinct prefronto-subcortical pathways in macaque monkeys, in other words, the pathways from PFC to CD (PFC-CD) and from PFC to MD (PFC-MD), we applied one of the chemogenetic tools, designer receptors exclusively activated by designer drugs (DREADDs). While neuronal silencing can be achieved by activating an inhibitory DREADD (hM4Di) following systemic agonist delivery (24), activating hM4Di expressed at axon terminals through local agonist infusion also leads to suppression of synaptic transmission (25–29). Here we developed imaging-guided chemogenetic synaptic silencing — a methodology for silencing neural pathways selectively with an agonist infused at hM4Di-positive PFC projection sites that are mapped in vivo beforehand by using positron emission tomography (PET) (30). Using this technique, we sequentially disconnected the bilateral/unilateral PFC-CD and PFC-MD pathways reversibly to investigate the causal roles in working memory and decision-making, which we hypothesized are distinctively governed by the two prefronto-subcortical pathways.

## Results

### PET visualizes hM4Di-positive terminal sites in CD and MD in vivo

We tested two monkeys in which hM4Di had been introduced into the bilateral PFC around the principal sulcus (Brodmann’s area 46) through adeno-associated virus (AAV) vector injections (Fig. 1A), because this area has been shown to play central roles in higher-order executive functions, including working memory and decision-making (1, 31). Several weeks after the injections, we performed PET imaging with a radiolabeled form of the highly-potent DREADD-specific agonist, deschloroclozapine (DCZ) (30), to visualize hM4Di expression non-invasively. Increased [11C]DCZ binding was localized not only in regions of the bilateral PFC (Fig. 1B), but was also visible as spots with a diameter of 3-5 mm in the CD head and lateral MD (Figs. 1C,D). These PET signals reflected hM4Di expression at both the soma/dendrite and the axon terminal sites, confirmed by immunohistochemistry in Mk1 (Fig. S1) and another monkey (Mk3) that received injections of an AAV vector encoding a fluorescent marker (monomergic Kusabira Orange, mKO) similarly into the PFC (Figs. 1E-J). Thus, we non-invasively identified the target sites for local agonist infusion to reversibly disconnect either the PFC-CD or PFC-MD pathway individually.

**Fig. 1.**
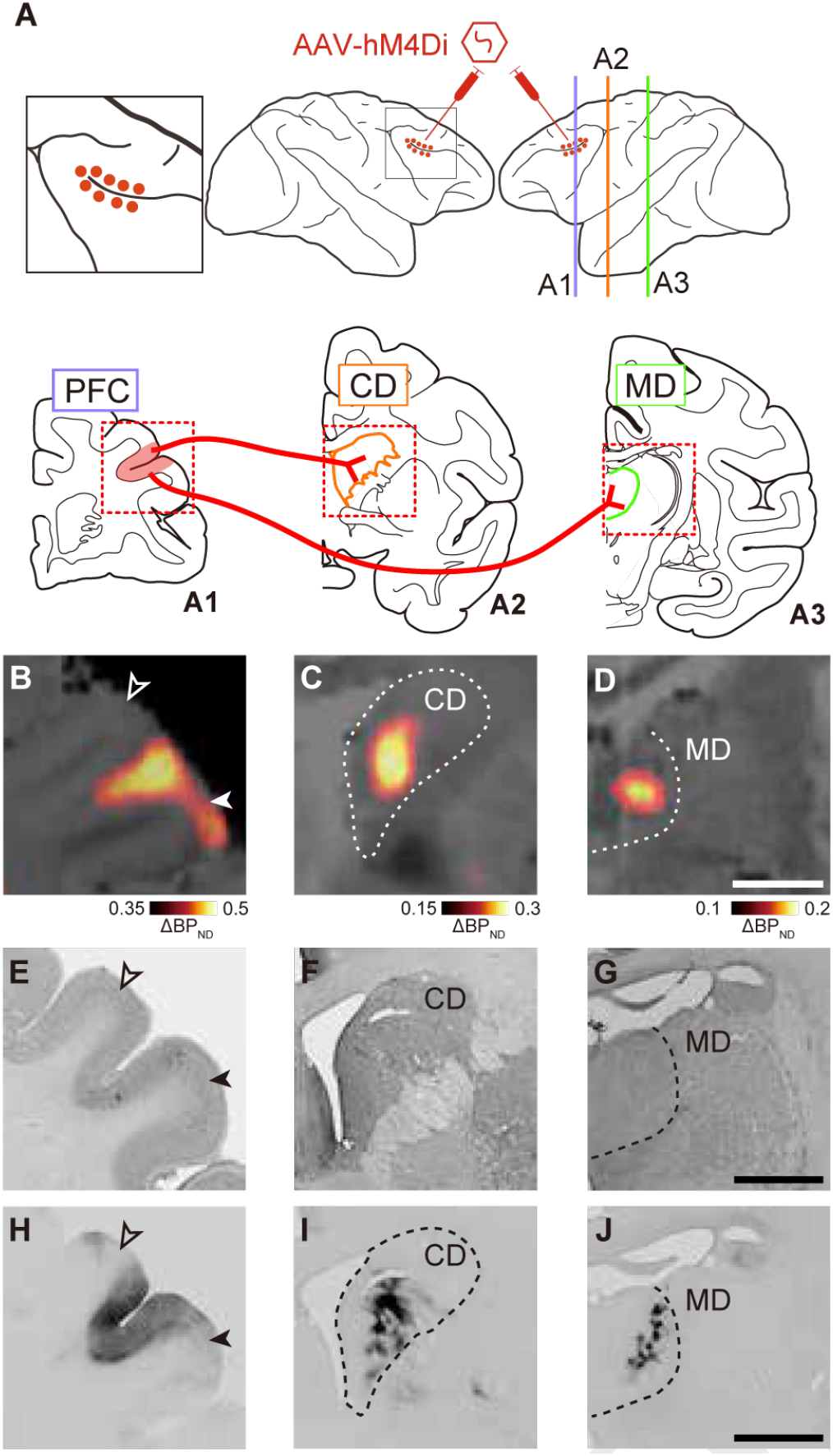
PET visualizes hM4Di-positive terminal sites in CD and MD in vivo. (A) Illustration representing the target pathways for chemogenetic silencing. Upper: Lateral view of injection sites. The left panel represents a zoomed-in view of the framed area of the center panel. Lower: Coronal planes corresponding to the lines in the lateral view. Red dots (upper) and shaded area (lower) represent intended injection sites (PFC) of an AAV vector expressing hM4Di, while red wires represent projections from PFC to CD and MD. (B-D) In vivo visualization of hM4Di expression in PFC (B), CD (C), and MD (D). Coronal PET image showing specific binding of [11C]DCZ subtracted before, from after the introduction of hM4Di, overlaying MR image of Mk2. (E-G) Corresponding nissl-stained sections in PFC (E), CD (F), and MD (G), respectively. (H-J) Corresponding DAB-stained sections representing immunoreactivity against a reporter protein (mKO) in PFC (H), CD (I), and MD (J) of a reference monkey that received viral vector injections into PFC in a similar manner. Open and filled arrowheads represent the dorsal and ventral b orders of the target regions, respectively. Scale bars: 5 mm.

### Chemogenetic silencing of the bilateral PFC-MD pathways selectively impaired the performance of a delayed response task

We then explored the significance of the PFC-subcortical pathways in working memory function in the monkeys who performed a spatial delayed response task (Fig. 2A). Prior to pathway-selective silencing, we verified the effect of chemogenetic silencing of the bilateral PFC. Consistent with our previous findings (30), systemic delivery of a low dose of DCZ (100 μg/kg, i.m.) drastically impaired the task performance in a delay-dependent manner, indicating that chemogenetic silencing of the bilateral PFC caused working memory deficits (Fig. 2B,E). Subsequently, we attempted to silence the PFC terminals through microinfusion of DCZ solution into the hM4Di-positive regions of either MD or CD bilaterally (Fig. 2C,D, upper). Infusion cannulae were placed under the guidance of CT images overlaying PET and MR images (Fig. 2C,D, lower). We adopted 100 nM of DCZ solution for microinfusion, because 1) preceding studies have used CNO (clozapine-N-oxide, the most frequently used DREADDs agonist) for local administration at 3 μM - 1 mM concentration (25, 26), which corresponds to 30 nM - 100 μM of DCZ in terms of its 100-fold higher agonist potency for hM4Di, and 2) 100 nM of DCZ is low enough to avoid off-target effects considering its affinities for numerous CNS receptors or transporters (30). In both monkeys, local DCZ infusion into MD prominently impaired the task performance, as compared to the control vehicle infusion (Fig. 2G). Impaired behavior was consistently observed within a single testing session (30 to 90 min after infusion; Fig. S2) and across the sessions (treatment × session, F(4,8) = 2.7, p = 0.11). Such impairment also depended on the delay duration (Fig. 2G), indicating that the PFC-MD pathway is critical for working memory function. By contrast, DCZ infusion into CD had no significant effect on the task performance (Fig. 2F).

**Fig. 2.**
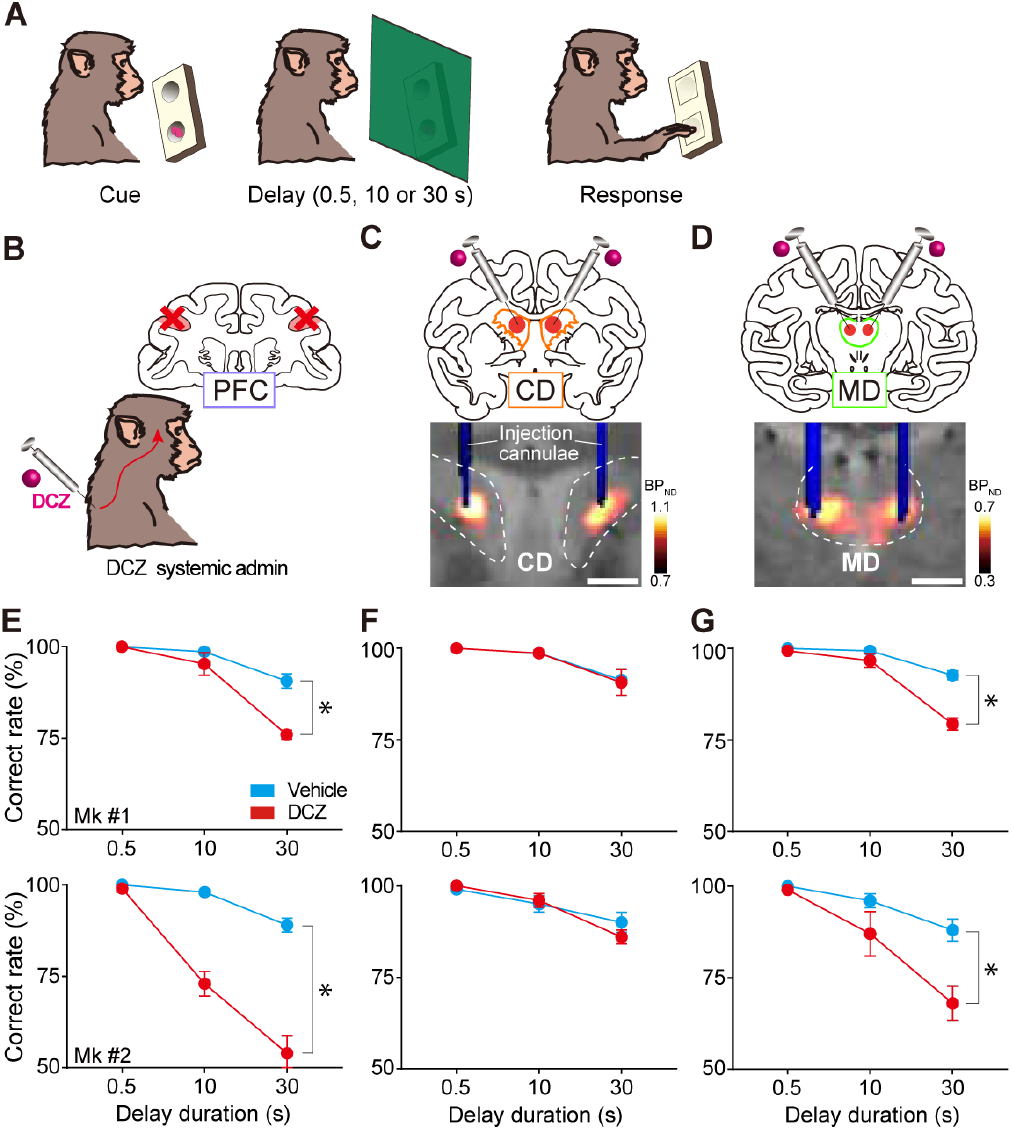
Chemogenetic silencing of the bilateral PFC-MD pathway selectively impaired the performance of a delayed response task. (A) Delayed response task. (B) Schematic diagram representing inactivation of PFC by systemic DCZ administration. (C) Chemogenetic silencing of the PFC-CD pathway by local DCZ infusion (100 nM, 3 μL/site) into bilateral CD, specifically a t h M4Di-positive P FC terminals sites. A CT image showing the infusion cannulae (blue) overlaying a structural MR image (gray), and a PET image showing increased [11C]DCZ binding (hM4Di expression, hot color). (D) Chemogenetic silencing of the PFC-MD pathway by local DCZ infusion (2 μL/site) into bilateral MD. (E) Behavioral effects of chemogenetic PFC inactivation. Correct performance rate (mean ± sem, n = 5) following DCZ (red) and vehicle infusions (cyan) are shown for two monkeys (two-way ANOVA with treatment × delay, main effect of treatment; Mk1, F(1,24) = 20.8, p = 1.2 × 10-4; Mk2, F(1,24) = 90.8, p =1.3 × 10-9). (F) Same as (E) but for the PFC-CD pathway (treatment; Mk1, F(1,24) = 0.03, p = 0.86; Mk2, F(1,24) = 0.20, p = 0.66). (G) Same as (E) but for the PFC-MD pathway (treatment; Mk1, F(1,24) = 32.9, p = 6.6 × 10-6; MK2, F(1,24) = 12.6, p = 1.6 × 10-3; treatment × delay; Mk1, F(2,24) = 16.5, p = 3.1 × 10-5; Mk2, F(2,24) = 3.8, p = 3.6 × 10-2). Asterisks: p < 0.05. Scale bars: 5 mm.

### Chemogenetic silencing of the unilateral PFC-CD pathway altered the choice preference in a delayed response task

Having demonstrated effective bilateral pathway-specific silencing, we next examined the effects of unilateral silencing of each pathway on the task in the same animals. We expected that unilateral silencing would cause an imbalance in the information flow, thereby resulting in asymmetric behavioral changes between hemifields. We infused DCZ solution into either CD or MD (3 μL and 2 μL, respectively), which would not diffuse beyond the midline, warranting unilateral pathway-specific silencing (see STAR Methods). In contrast to the bilateral silencing, local DCZ infusion into unilateral MD did not affect the performance of delayed response task in either monkey (Figs. 3F,G), suggesting that intact function of either PFC-MD pathway is sufficient for cognitive behavior as was implicated in previous lesion studies in monkeys (32). Local DCZ infusion into unilateral CD, on the other hand, significantly decreased the correct performance rate in trials with food given in the contralateral well to the injected side (Fig. 3E), and tended to improve the performance on the ipsilateral side (Fig. 3D). Given that the bilateral silencing spared working memory function, it was unlikely to be due to a loss of working memory. Together, these results suggest that the PFC-MD pathway, but not PFC-CD pathway, plays an essential role in working memory function, where both sides of the pathways can complement each other.

**Fig. 3.**
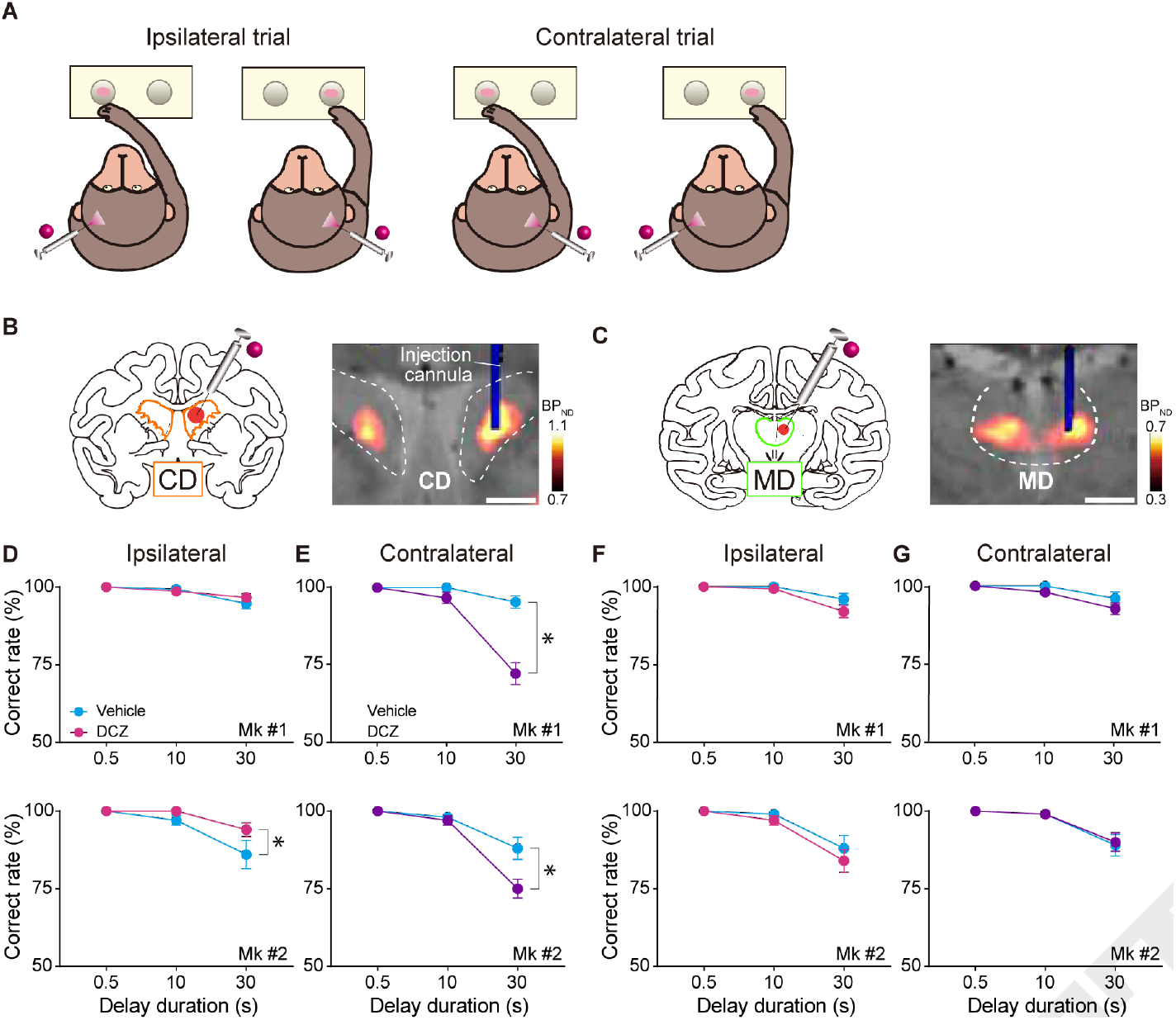
Chemogenetic silencing of the unilateral PFC-CD pathway altered the choice preference in a delayed response task. (A) Illustrations representing trial types (ipsilateral and contralateral) of the delayed response task. (B,C) Chemogenetic silencing of the PFC-CD (B) and PFC-MD (C) pathways by local DCZ infusion into unilateral CD (3 μL/site) and MD (2 μL/site), respectively. CT image showing the infusion cannula (blue) overlaying on MR image (anatomy, gray) and PET image showing increased [11C]DCZ binding (hM4Di expression, hot color). (D,E) Behavioral effects of chemogenetic silencing of the unilateral PFC-CD pathway for ipsilateral (D) and contralateral (E) trials (ipsilateral; Mk1, F(1,54) = 0.34, p = 0.56; Mk2, F(1,54) = 4.4, p = 4.1 × 10-2; contralateral; Mk1, F(1,54) = 35.8, p = 1.7 × 10-7; Mk2, F(1,54) = 7.4, p = 8.7 × 10-3). (F,G) Same as (D,E) but for silencing of the unilateral PFC-MD pathway (ipsilateral; Mk1, F(1,54) = 3.0, p = 0.09; Mk2, F(1,54) = 1.0, p = 0.31; contralateral; Mk1, F(1,54) = 3.3, p = 0.07; Mk2, F(1,54) = 0.04, p = 0.84). Correct performance rate (mean ± sem, total 10 sessions, 5 left and 5 right infusions for each target) following DCZ (ipsilateral, magenta; contralateral, purple) and vehicle infusions (cyan) are shown for two monkeys. Asterisks: p < 0.05. Scale bars: 5 mm.

### Chemogenetic silencing of the PFC-CD pathway induced laterality bias in a free-choice task

As demonstrated above, DCZ infusion into unilateral CD led to hemifield-dependent behavioral changes in a working memory performance, while the bilateral injections did not. These observations led us to postulate that unilateral silencing of the PFC-CD pathway induces laterality bias in decision-making in a hemifield-dependent fashion as implicated in previous single-unit recording and lesion studies showing an involvement of CD in such a phenomenon (33). To identify the contribution of the PFC-CD/MD pathways in decision-making — another function being responsible for the prefronto-subcortical network, we focused on a behavioral bias under a free-choice situation. We used a free-choice task, in which the monkeys were allowed to pick up food from either left or right baited wells without instructions (Fig. 4A). If signals processed via the PFC-CD pathway promoted contralateral choices, silencing of this pathway on one side of the brain would increase a choice of wells on the same side. As we predicted, unilateral DCZ infusion into CD (Fig. 4B) significantly increased the choice of ipsilateral wells compared with the vehicle infusion, indicating that ipsilateral bias emerged (Fig. 4D). This impairment was not due to the inability of voluntary attention to the contra-hemifield (i.e., contralateral hemineglect) as seen in the case of unilateral dopamine depletion from CD (34), because the animals maintained normal spontaneous eye movements by viewing both the ipsi- and contra-hemifields at the same frequency (Fig. S3). Unilateral DCZ infusion into MD (Fig. 4C), on the other hand, did not change the tendency of the monkeys’ choices (Fig. 4E). Thus, the overall results suggest that the PFC-CD pathway in each hemisphere may contribute to decision-making by biasing one’s action toward the contralateral side without direct involvement of working memory or oculomotor function.

**Fig. 4.**
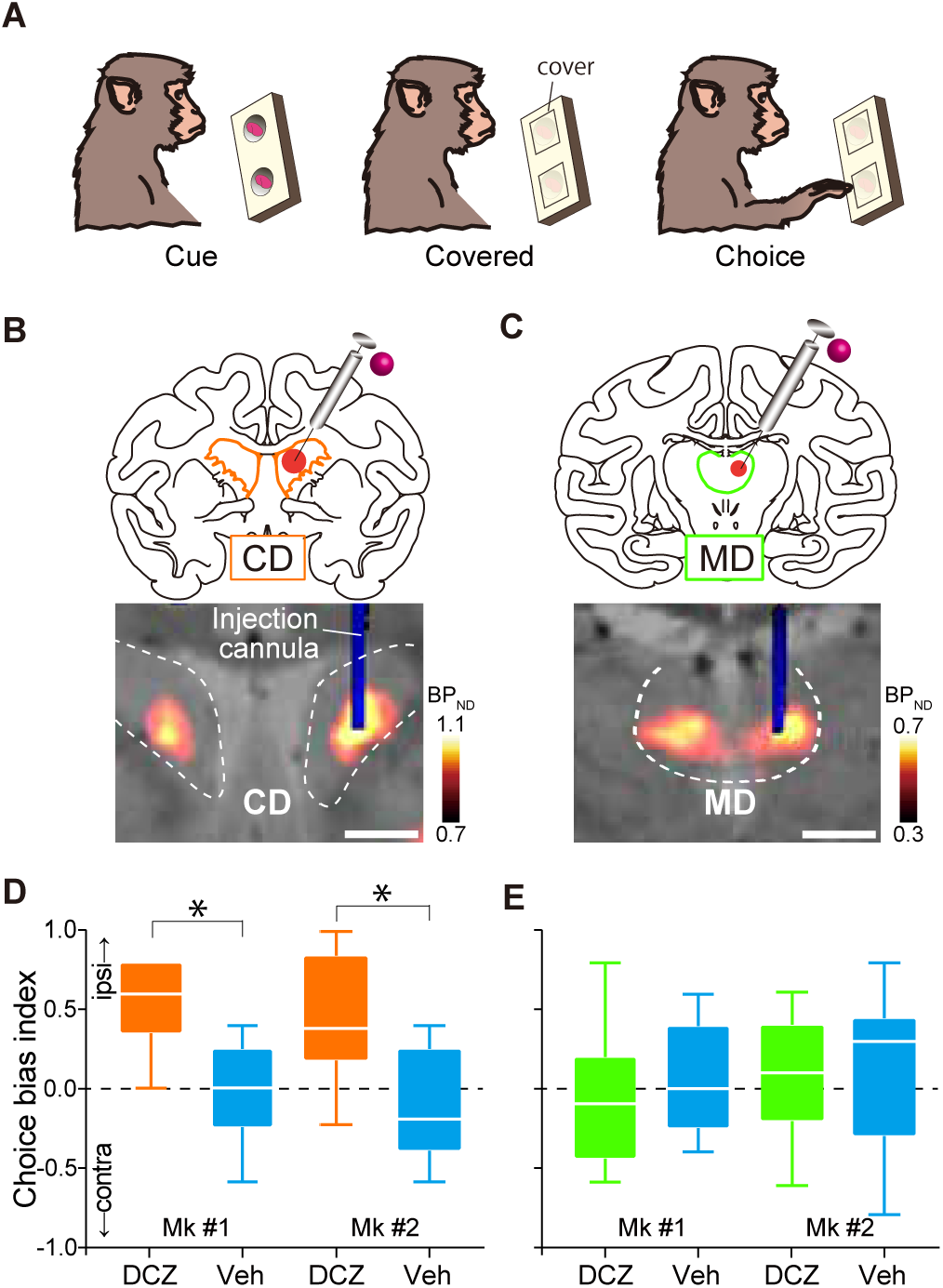
Chemogenetic silencing of the unilateral PFC-CD pathway induced laterality bias in a free-choice task. (A) Free choice task. (B,C) Chemogenetic unilateral silencing of the PFC-CD (B) and PFC-MD pathways (C) by local unilateral DCZ infusion. Arrangements are the same as in Fig. 2C,D. (D,E) Choice bias index of free-choice task following unilateral DCZ infusion into CD (D, orange; Brunner-Munzel Test; Mk1, BM-value = −6.7, p = 3.3 × 10-6; Mk2, BM-value = −5.0, p = 9.4 × 10-5) and MD (E, green; Mk1, BM-value = 0.79, p = 0.44; Mk2, BM-value = 0.37, p = 0.72), and control vehicle infusions (cyan). Boxplots indicate the median (line within the box), the 1st–3rd quartiles (box), and the largest and smallest data points (whiskers) (total 10 sessions, 5 left and 5 right infusions for each target). Asterisks: p < 0.05. Scale bars: 5 mm.

Finally, we verified that any behavioral side-effects were not caused by the effect of DCZ infusion into non-target sites, i.e., areas without innervations of hM4Di-positive axons. We infused DCZ into brain regions where significant hM4Di expression was undetected, including the ventral putamen and ventroanterior thalamus, the structures located adjacent to CD and MD, respectively. In these cases, the monkeys displayed no significant behavioral changes in either task (Fig. S4), thus supporting that our data described above are attributable to the chemogenetic silencing of hM4Di-positive PFC-derived axon terminals.

## Discussion

Here we chemogenetically dissected the neuronal information flow from the PFC to two subcortical areas, MD and CD, in individual monkeys. Our results provided direct evidence for a functional dissociation between PFC-MD and PFC-CD pathways on the performance of the delayed response task and free-choice task. We also showed that bilateral and unilateral silencing had different impacts on behavior, with their effects being consistent across tasks. Our present study is the first, to our knowledge, to demonstrate a dissociation of functional roles of these two key prefronto-subcortical pathways. Although the detailed mechanism remained to be identified, our results would provide new insights into the neural basis for higher-order executive functions.

Anatomically, PFC and CD/MD constitute two loop circuits, namely the cortico-basal ganglia-thalamo-cortical loop and the cortico-thalamo-cortical loop. The former is a part of a series of parallel-projecting, largely segregated circuits that arise from the cortex and link the striatum (including CD), pallidum, and thalamus back to the same original cortical area (14, 35). The latter is a reciprocal connection between the cortex and thalamus (36, 37). Thus, PFC communicates with two partially overlapped subcortical circuits, where MD receives information from the same PFC area both directly and indirectly. Unlike conventional inactivation or lesion experiments, the chemogenetic manipulation described here dissected the PFC to the CD/MD pathways independently, thereby allowing us to untangle the direct and indirect effects on MD. Our findings suggest that the PFC-subcortical projections originating in the same region play distinct roles in cognitive functions. What is the mechanistic explanation for the differential contribution? One possibility is that the two pathways originated from distinct neuronal populations that distinctly signal critical information for working-memory or decision-making. It has been shown that the majority of both PFC-MD and PFC-CD projections arise from deep layers (12, 13), while a small proportion of PFC neurons have axonal branches that innervate both sites (38). Thus there might exist anatomically intermingled subpopulations in the PFC. Another possibility is that the two pathways convey similar information, which is processed differently in each subcortical area, leading to the distinct functions. Further study would be required to clarify the underlying mechanism of the causal role of the two pathways. Whatever the mechanism is, our results indicate that a single PFC region exerts different cognitive functions depending on which subcortical area it interacts with. These pathway-specific functions have been overlooked, because they cannot be distinguished in conventional neuronal recording or inactivation experiments. Indeed, the results of these experiments have demonstrated functional localization in multiple PFC areas (1). Our results extend this view and highlight the functional heterogeneity of PFC-subcortical circuits in higher-order executive functions.

The importance of PFC and MD in implementing working memory has been repeatedly shown (8, 39). A human imaging study reported that the strength of functional connectivity between PFC and MD during a working memory task correlated with memory load (11), suggesting their cooperative involvement. In a monkey study, similar proportions of neurons in PFC and MD were active during the delay period in working memory task, whereas a greater proportion of neurons in MD than PFC participated in the response phase, suggesting that the PFC-MD pathway is involved in both maintenance and transformation of working memory (40). Our findings corroborated these notions by providing direct evidence for the causal involvement of the primate PFC-MD pathway in working memory. Our results also extend the recent findings in a rodent study that showed an essential role of the PFC-MD pathway in a spatial working memory task (41), and together imply possible common neural substrates between primates and rodents.

Interestingly, silencing of the unilateral PFC-MD pathway did not have an impact on the behavioral performance of the delayed response task. This result suggests that the unilateral dysfunction of this pathway would be compensated by the other, which contrasted with the case in which the unilateral dysfunction of PFC per se that has been reported to cause deficits in contra-hemifield working memory (42). This finding also raised the possibility that the PFC-MD pathway would convey non-lateralized information in terms of visuospatial processing, at least in relation to implementing working memory, which was supported by a previous electrophysiological recording in monkey MD (43). Future studies that combine electrophysiological recording and pathway-specific silencing would be required to reveal the essential information flow from PFC to MD.

On the other hand, our findings also suggest that the PFC-CD pathway does not play an essential role in the performance of a spatial delayed response task, which is one of the major paradigms when examining spatial working memory (44, 45). However, this does not necessarily mean that the PFC-CD pathway is irrelevant for working memory. It has been proposed that PFC-striatal interactions act as a gate for selecting the contents of working memory that should be stored through the prefronto-basal ganglia-thalamus network (46, 47). For example, functional connectivity between the dorsolateral part of the PFC to the anterior striatum in human (presumably homologous to PFC-CD in monkeys) was modulated by the novelty of visual cues to be memorized (48). In our task setting, monkeys were just required to memorize the location of the familiar foods, which might not require contribution of the PFC-CD pathway due to its simplicity. Taken together, our results show that the PFC-CD pathway might not play a pivotal role in implementing working memory, but they do not exclude the possibility that this pathway contributes to supporting working memory in a more complicated situation.

Another key finding of our study is that unilateral silencing of the PFC-CD pathway resulted in biasing choices away from the contralateral option under the free-choice situation (Fig. 4). Unlike the visual hemineglect induced by unilateral dopamine depletion in CD (34, 49), we did not find any significant deviation of spontaneous eye position (Fig. S3). Thus the deficits observed here were not directly caused by alteration of eye-movement or involuntary attention. Similar ipsiversive bias in a free-choice task has been reported to occur after unilateral deactivation of the dorsolateral part of the PFC in monkeys (50). It has been proposed that the pathway from the PFC to CD conveys cognitive information while CD neurons represent specific actions/stimuli associated with value preferentially in the contralateral hemisphere to guide goal-directed behavior (51, 52). We speculate that unilateral silencing of the PFC-CD pathway may disrupt the contra-preferred value representation in CD, leading to an increase in ipsilateral choices. This also explains our paradoxical result that unilateral, but not bilateral, PFC-CD silencing produced behavioral bias in the delayed response task. Contrary to the PFC-CD pathway, unilateral silencing of the PFC-MD pathway did not alter the performance of the free-choice task. Because the primate MD is known to be involved in adaptive decision-making, such as value-based choice behavior (8), our results are possibly attributable to compensation by the intact side of the PFC-MD pathway that may process non-lateralized information, as discussed above.

Recent advances in specific neuronal manipulation tools have drastically enhanced our understandings of the functional roles of individual neural pathways in small animals such as rodents (24, 53). Nonetheless, the application of pathway-selective manipulations to non-human primates is still largely limited by using either optogenetic terminal activation (19–21) or double-vector methods (15–18), neither of which is capable of testing dissociable function of multiple neural pathways in individual subjects. Using a chemogenetic approach, the current study succeeded in the double-dissociation of the roles of two distinct pathways in individual monkeys. Our findings were largely owing to highly reproducible manipulations of a specific pathway, which was achieved by in vivo projection mapping of the hM4Di-positive sites in CD/MD and accurate positioning of the injection cannulae into the areas proven by CT imaging following every injection. The imaging-guided chemogenetic synaptic silencing permits us to identify and dissect pre-defined circuits. In combination with methods to monitor neuronal activity, such as electrophysiological recording or functional neuroimaging, it would be possible to specify the precise content of signals processed through the prefronto-subcortical network. Given the accurate targeting with applicable capability, imaging-guided chemogenetics affords powerful and unique strategies for pathway-selective dissection, thereby expanding our opportunities of linking neural circuits to behaviors in primates.

In summary, the present study has demonstrated a dissociation of the cognitive roles that the two PFC pathways toward MD and CD exert in working memory and decision-making, respectively. Given the anatomical and functional heterogeneity of the primate PFC, and especially its dorsolateral part on which we focused (54), our findings provide invaluable insights toward the understanding of a neural basis for higher-order executive functions/dysfunctions associated with neuropsychiatric disorders.

## ACKNOWLEDGEMENTS

We thank J. Kamei, R. Yamaguchi, Y. Matsuda, Y. Sugii, T. Okauchi, R. Suma, A. Maruyama, T. Kokufuta, Y. Iwasawa, T. Watanabe, A. Tanizawa, S. Shibata, N. Nitta, Y. Ozawa, M. Fujiwara, and M. Nakano for their technical assistance. Funding: This study was supported by MEXT/JSPS KAKENHI Grant Numbers JP18K15353 (to KO), JP17H02219 (to TH), JP19H05467 (to MT), JP15H05917 and JP18H04037 (to TM), by AMED Grant Numbers JP20dm0307007 (to TH), JP20dm0307021(to KI), JP18dm0207003 (to MT), JP20dm0107146 (to TM), by JST PRESTO Grant Number JPMJPR1683 (to KI), by the cooperative research program at PRI, Kyoto Univ., and by National Bio-Resource Project “Japanese Monkeys” of MEXT, Japan. Author contributions: Conceptualization, T.M.; Investigation, K.O., Y.H., and Y.N.; Resources, K.I. and M.T.; Visualization, K.O.; Writing – review editing, All authors. Competing interests: The authors declare no competing interests. Data and materials availability: The gene constructs of viral vectors and data sets used in this study are available from the corresponding author on request.

## Materials & Methods

### Subjects

All experimental procedures involving animals were carried out in accordance with the Guide for the Care and Use of Laboratory Animals (National Research Council of the US National Academy of Sciences) and were approved by the Animal Ethics Committee of the National Institutes for Quantum and Radiological Science and Technology. Three macaque monkeys [Mk1: male Rhesus (Macaca mulatta), Mk2: female Japanese monkey (Macaca fuscata), and Mk3: female Rhesus (Macaca mulatta); 5.0, 5.7, and 4.2 kg; age 5, 6 and 9 years at the beginning of experiments] were used. The monkeys were kept in individual primate cages in an air-conditioned room. A standard diet, supplementary fruits/vegetables and a tablet of vitamin C (200 mg) were provided daily.

### Viral vector production

AAV1 (AAV1-hSyn-hM4Di-IRES-AcGFP) and AAV2 (AAV2-CMV-mKO) vectors were produced by helper-free triple transfection procedure, which was purified by affinity chromatography (GE Healthcare, Chicago, USA). Viral titer was determined by quantitative PCR using Taq-Man technology (Life Technologies, Waltham, USA).

### Surgical procedures and viral vector injections

Surgeries were performed under aseptic conditions in a fully equipped operating suite. We monitored body temperature, heart rate, SpO2 and tidal CO2 throughout all surgical procedures. Monkeys were immobilized by intramuscular (i.m.) injection of ketamine (5-10 mg/kg) and xylazine (0.2-0.5 mg/kg) and intubated with an endotracheal tube. Anesthesia was maintained with isoflurane (1-3 %, to effect). Prior to surgery, magnetic resonance (MR) imaging (7 tesla 400mm/SS system, NIRS/KOBELCO/Brucker) and X-ray computed tomography (CT) scans (Accuitomo170, J. MORITA CO., Kyoto, Japan) were performed under anesthesia (continuous infusion of propofol 0.2-0.6 mg/kg/min, intravenously). Overlaid MR and CT images were created using PMOD® image analysis software (PMOD Technologies Ltd, Zurich, Switzerland) to estimate stereotaxic coordinates of target brain structures.

Two monkeys (Mks 1 and 2) were injected AAV1-hSyn-hM4Di-IRES-AcGFP (4.7 × 10e13 particles/mL) into the bilateral prefrontal cortex (Brodmann’s area 46), and one monkey (Mk3) was injected AAV2-CMV-mKO (1.7 × 10e13 particles/mL) into the unilateral prefrontal cortex. After retracting skin and galea, the frontal cortex was exposed by removing a bone flap and reflecting the dura mater. Handheld injections were made under visual guidance through an operating microscope (Leica M220, Leica Microsystems GmbH, Wetzlar, Germany), with care taken to place the beveled tip of the Hamilton syringe containing the viral vector at an oblique angle to the brain surface. The needle was inserted into the intended area of injection by one person and a second person pressed the plunger to expel approximately 1 μL. Nine tracks were injected in each hemisphere; one was located 1 mm posterior to the caudal tip of the principal sulcus, and the others were located along the dorsal (4 tracks) and ventral (4 tracks) bank of the principal sulcus posterior to the rostral tip of the ascending limb of the arcuate sulcus (Fig. 1). Viral vectors were injected 3 to 5 μL per track depending on the depth. Totals of 35 and 37 μL for Mk1 and 40 and 44 μL for Mk2 of viral aliquots were injected into the right and left hemispheres, respectively, and 20 μL into the left hemisphere of Mk3.

### Behavioral task

Two monkeys (Mks 1 and 2) were tested with a spatial delayed response task (Fig. 2). The protocol was almost the same as that of our previous study (30). Behavioral testing was conducted in a sound-attenuated room. Monkeys were seated in a monkey chair from which they could reach out one hand and take food to their mouths. A wooden table with two food-wells was placed in front of the monkeys, and a screen was placed between the monkeys and the table. First, a piece of food reward (raisin or snack) was placed in one of the two food-wells, and then both wells were covered with wooden plates. Then, the screen was placed for 0.5, 10 or 30 s, which served as delay periods. The position of the baited well (left or right) and the delay period (0.5, 10 or 30 s) were determined pseudo-randomly. After the delay period, the screen was removed, and the monkeys were allowed to select either food-well to get the food. The monkeys were allowed to get the food if they reached for the correct food-well and removed the cover plate. The inter-trial interval was set at 10 s. A daily session lasted about one hour, consisting of 3 blocks of 30 trials (10 trials for each delay of 0.5, 10, or 30 sec) for Mk1 and 2 blocks of 30 trials for Mk2. Each of the monkeys performed the single sessions individually. The blocks were separated by 5-min rest periods. The task was tested under both bilateral and unilateral silencing conditions.

Two monkeys (Mks 1 and 2) were also tested with a free choice task only under unilateral silencing conditions (Fig. 4). In this task, both wells were baited, and then the food wells were covered. Then, the table was placed in front of the monkeys and they were allowed to get the food. After the monkeys picked up one side of food, the food table was withdrawn, and then both wells were baited again for the next trial. The task consisted of 10 trials. The delayed response task and the free choice task were conducted in the same sessions in a counterbalanced sequence.

### Drug administration

For systemic i.m. injection, DCZ was dissolved in 2% of dimethyl sulfoxide (DMSO)/saline to a final volume of 0.1 mg/kg and injected intramuscularly 15 min before the beginning of the experiments. Note that DCZ itself did not affect the performance in monkeys without hM4Di expression (30, 55). For microinfusion, DCZ was first dissolved in DMSO and then diluted in PBS to a final concentration of 100 n M. We prepared fresh solutions on the day of usage. We used two stainless steel infusion cannulae (outer diameter 300 μm; Muromachi-Kikai) inserted into each target region: caudate nucleus (CD) and mediodorsal thalamus (MD), and ventroanterior thalamus and ventral putamen for controls. Each cannula was connected to a 10-μL microsyringe (7105KH; Hamilton) via polyethylene tubing. These cannulae were advanced through the guide tube by means of an oil-drive micromanipulator. DCZ or PBS was injected at a rate of 0.25 μL/min by auto-injector (Legato210; KD Scientific) for a total volume of 3 μL for CD and putamen and 2 μL for MD and ventroanterior thalamus for each hemisphere. For unilateral silencing, we injected either DCZ or PBS into one hemisphere. The injection volumes were determined based on a previous study reporting that injection of 3 μL and 1.5 μL aqueous had spread about 5–6 mm and 3–4 mm in diameter in monkey brain, respectively (56). We chose enough volumes (3 μL and 2 μL for CD and MD, respectively) to cover each hM4Di-positive terminal site as we observed clear PET signals with a diameter of 5–7 mm and 3–4 mm for CD and MD in each hemisphere, respectively. Importantly, the cannula(e) was placed at least 4-mm apart from the midline for both targets (CD or MD), so that the injected solution would not diffuse beyond the midline, warranting unilateral silencing of a specific pathway when injected unilaterally. The behavioral session began about 30 min after the infusion was finished and lasted about one hour. We performed at most one silencing experiment per week for one area. At the end of each infusion, a CT image was obtained to visualize the infusion cannulae in relation to the chambers and skull. The CT image was overlaid on MR and PET images by using PMOD to verify that the infusion sites (tips of the infusion cannulae) were located in the areas in which differential PET signals were observed when radioactive DCZ was administered, indicating that the area presumably received projections from DLPFC. As a control experiment, we also injected DCZ solution into the ventroanterior, located anterior to MD, or putamen, where significant hM4Di expression was not observed by PET (Fig. S4).

Following several unilateral infusions of DCZ into CD, we monitored the monkeys’ eye movements while their head was fixed b y u sing an eye-tracking camera (ETL-200, ISCAN) located 40 cm from the monkeys’ eyes. Data were stored on a computer and then processed by custom-made software on Matlab (Mathworks).

### PET imaging

PET imaging was conducted as previously reported (30). Briefly, PET scans were conducted at 45 days after injection of vectors for both monkeys, and also before injection of vectors for Mk2. PET scans were performed using a microPET Focus 220 scanner (Siemens Medical Solutions USA, Malvern, USA). The monkeys were immobilized by ketamine (5-10 mg/kg) and xylazine (0.2-0.5 mg/kg) and then maintained under anesthetized condition with isoflurane (1-3%) during all P ET procedures. Transmission scans were performed for about 20 min with a Ge-68 source. Emission scans were acquired in 3D list mode with an energy window of 350–750 keV after an intravenous (i.v.) bolus injection of [11C]DCZ (344.8-369.6 MBq). Emission data acquisition lasted 90 min. PET image reconstruction was performed with filtered back-projection using a Hanning filter cut-off at a Nyquist frequency of 0.5 m m–1. To estimate the specific binding of [11C]DCZ, regional binding potential relative to nondisplaceable radioligand (BPND) was calculated by PMOD® with an original multilinear reference tissue model (MRTMo) using the cerebellum as a reference.

### Histology and immunostaining

For histological inspection, two monkeys (Mk1 and Mk3) were deeply anesthetized with an overdose of sodium pentobarbital (80 mg/kg, i.v.) and transcardially perfused with saline at 4°C, followed by 4% paraformaldehyde in 0.1 M phosphate buffered saline (PBS), pH 7.4. The brain was removed from the skull, postfixed in the same fresh fixative overnight, saturated with 30% sucrose in phosphate buffer (PB) at 4°C, and then cut serially into 50-μm-thick sections on a freezing microtome. For visualization of immunoreactive signals of GFP (co-expressed with hM4Di) and mKO in Mk1 and MK3, respectively, a series of every 6th section was immersed in 1% skim milk for 1 h at room temperature and incubated overnight at 4°C with rabbit anti-GFP monoclonal antibody (1:500, G10362, Thermo Fisher Scientific) and rabbit anti-mKO polyclonal antibody (1:500; PM051M, MEDICAL BIOLOGICAL LABORATORIES CO., LTD, Japan), respectively, in PBS containing 0.1% Triton X-100 and 1% normal goat serum for 2 days at 4°C. The sections were then incubated in the same fresh medium containing biotinylated goat anti-rabbit IgG antibody (1:1,000; Jackson ImmunoResearch, West Grove, PA, USA) for 2 h at room temperature, followed by avidin-biotin-peroxidase complex (ABC Elite, Vector Laboratories, Burlingame, CA, USA) for 2 h at room temperature. For visualization of the antigen, the sections were reacted in 0.05 M Tris-HCl buffer (pH 7.6) containing 0.04% diaminobenzidine (DAB), 0.04% NiCl2, and 0.003% H2O2. The sections were mounted on gelatin-coated glass slides, air-dried, and cover-slipped. A part of other sections was Nissl-stained with 1% Cresyl violet. Images of sections were digitally captured using an optical microscope equipped with a high-grade charge-coupled device (CCD) camera (Biorevo, Keyence, Osaka, Japan).

### Statistics

To examine the effect of each treatment on the performance of the delayed response task, behavioral measurement (correct rates) was subjected to two-way ANOVA (treatment × delay) using GraphPad Prism 7. For the free-choice task, the choice index was calculated by the following equation:

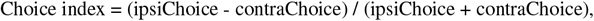

where ipsiChoice and contraChoice indicate the number of ipsilateral and contralateral choices, respectively, against a hemisphere in which DCZ solution was injected. Then the index was subjected to Brunner-Munzel test to compare with vehicle infusions using R.

## Supplementary Materials

**Figure S1.**
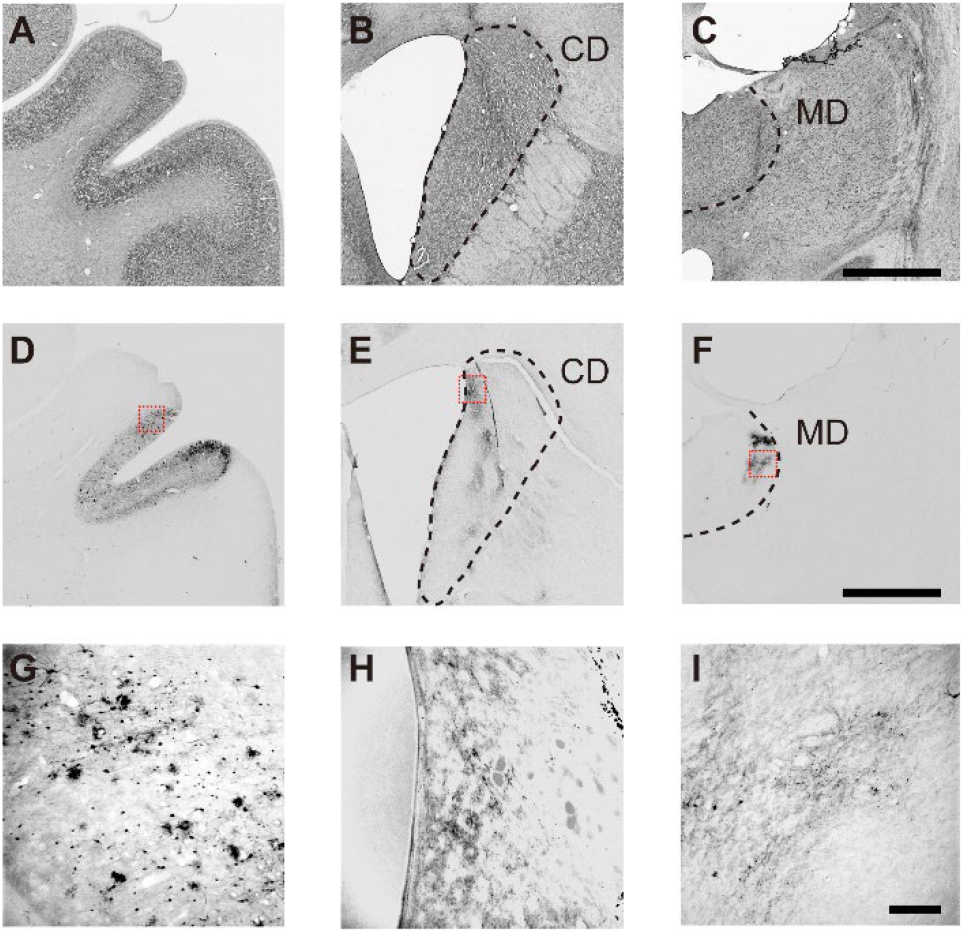
Histological sections of Mk#1 for identifying expression of hM4Di. (**A-C**) Nissl-stained sections in PFC (A), CD (B), and MD (C), respectively, corresponding to PET images in Fig. 1. (**D-F**) Corresponding DAB-stained sections representing immunoreactivity against a reporter protein (GFP) co-expressed with hM4Di in PFC (D), CD (E), and MD (F). (**G-I**) Zoomed-in views of red-squared areas in D-F, respectively. Scale bars: 5 mm for A-F and 250 μm for G-I.

**Figure S2.**
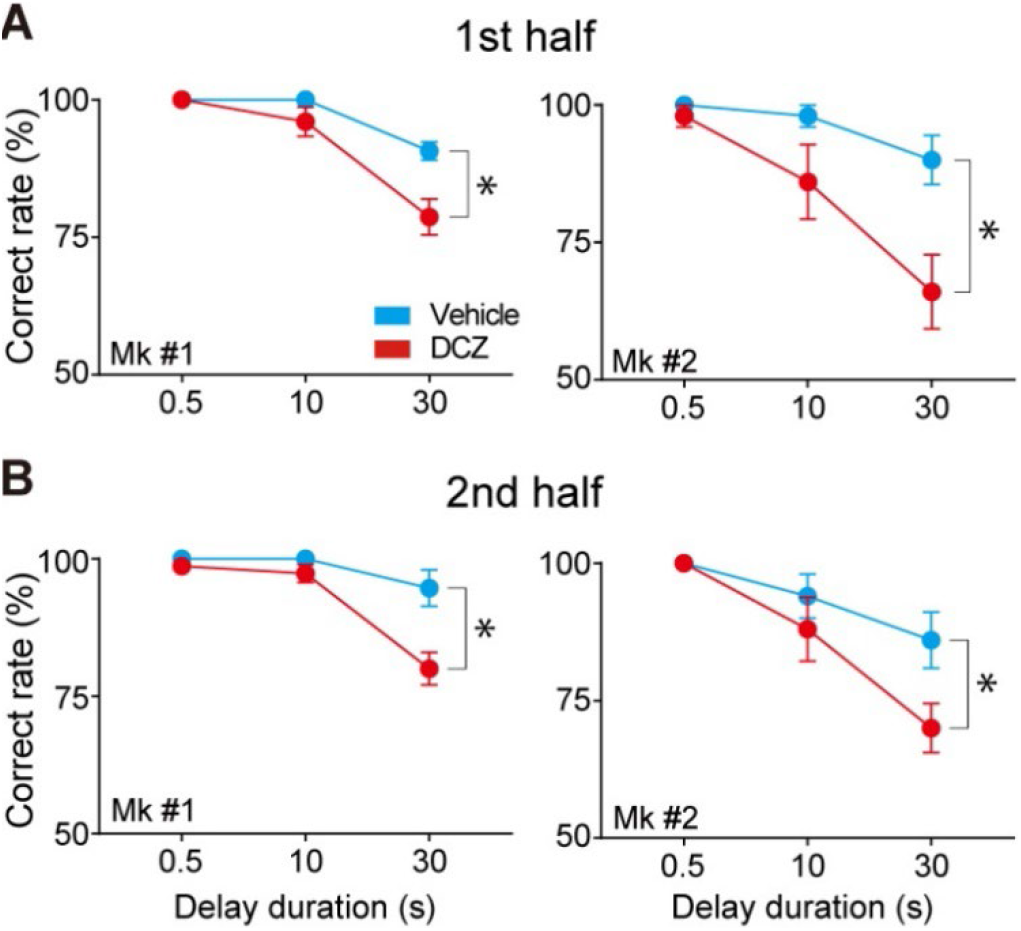
Consistent behavioral effects of DCZ infusion during early vs. late phases of session. (**A,B**) Comparison of behavioral effects of local DCZ infusion into bilateral MD in the 1st (A) and 2nd half (B) of a session, respectively. Correct performance rates (mean ± sem) following local infusion of DCZ (red) and control vehicle infusion (cyan) are shown for monkeys #1 (left) and #2 (right). Asterisks indicate that DCZ infusion had significant effects on performance compared to vehicle control for all cases (1st half, two-way ANOVA, main effect of treatment; Mk#1, F_(1,24)_ = 12.5, p = 1.7 × 10^−3^; Mk#2, F_(1,24)_ = 12.0, p = 2.0 × 10^−3^; 2nd half, Mk#1, F_(1,24)_ = 14.5, p = 8.5 × 10^−4^; Mk#2, F_(1,24)_ = 5.0, p = 3.4 × 10^−2^). Three-way ANOVA (treatment × delay × time) also revealed that there was no significant effect (Mk#1, F_(1,48)_ = 0.16, p = 0.69; Mk#2, F_(1,48)_ = 1.2, p = 0.26) in the interaction of treatment and time, suggesting that the effect of local DCZ infusion did not differ between the early and late phases of a session.

**Figure S3.**
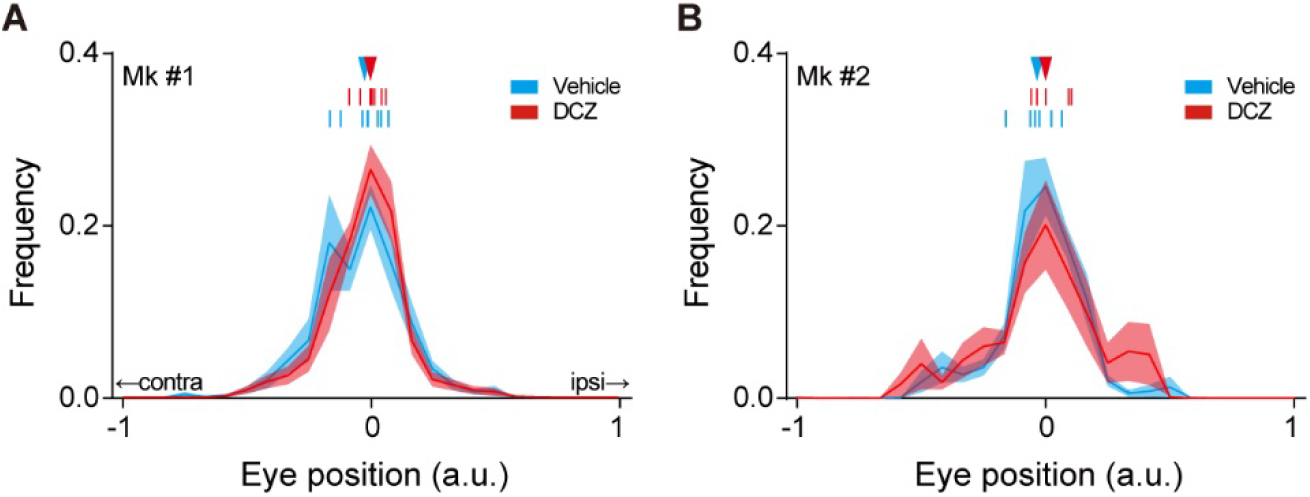
Effect of unilateral DCZ infusion into CD on spontaneous eye movements. (**A,B**) Histograms of the frequency of horizontal eye position from center (0) following local infusion of DCZ (red) and control vehicle infusion (cyan) into CD for Mks #1 (n = 8 sessions, 4 left and 4 right infusions into each side) and #2 (n = 6 sessions, 3 left and 3 right infusions), respectively. Solid lines and shaded areas represent mean and sem, respectively. For both monkeys, DCZ administration had no significant effects on averaged distribution of eye movements compared to vehicle control (Kolmogorov-Smirnov test; Mk#1, D = 0.12, p = 0.99; Mk#2, D = 0.24, p = 0.47). Ticks above the histograms represent the median of horizontal eye position in each session, and triangles represent those means, respectively. For both monkeys, DCZ administration had no significant effects on medians compared to vehicle control (Welch’s t-test; Mk#1, t_11.2_ = 0.17, p = 0.47; Mk#2, t_9.9_ = 1.0, p = 0.32).

**Figure S4.**
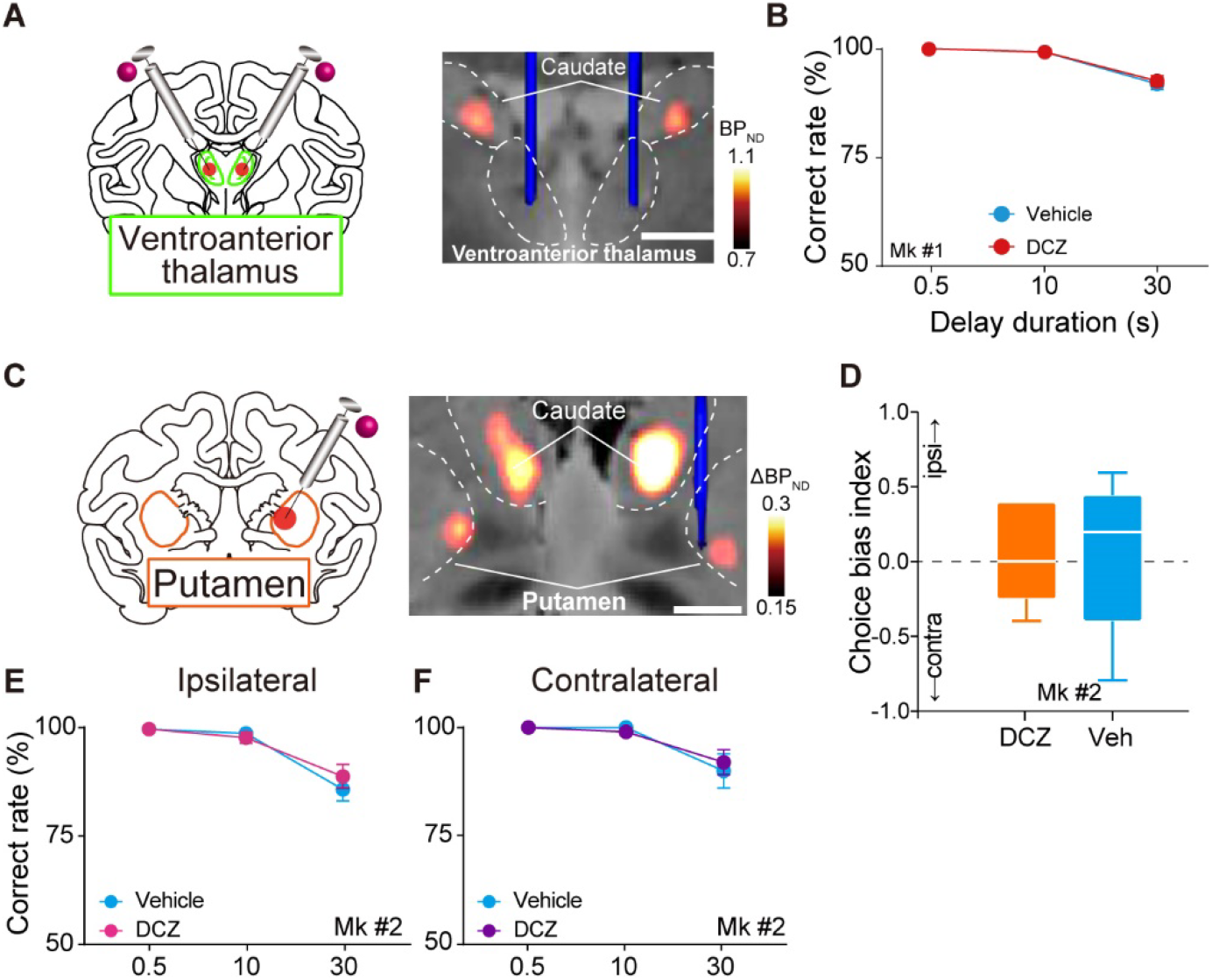
Effects of local infusion of DCZ into non-DREADD regions. (**A**) Local DCZ infusion into the bilateral ventroanterior thalamus (2 μL/site). CT image showing location of infusion cannulae (blue) overlaying MR image (anatomy, gray) and PET images showing increased [^11^C]DCZ binding (hM4Di expression, hot color). (**B**) Behavioral effects of local DCZ infusion into bilateral ventroanterior thalamus. Correct performance rates following local infusion of DCZ (red) and control vehicle infusion (cyan) are shown for one monkey. Local DCZ infusion into bilateral ventroanterior thalamus had no significant effect on performance (two-way ANOVA, main effect of treatment; F_(1,24)_ = 0.11, p = 0.75). (**C**) Local DCZ infusion into unilateral ventral putamen (3 μL/site). (**D**) Choice bias index of free-choice task following unilateral DCZ infusion into ventral putamen (orange) and vehicle infusion (cyan). Boxplot shows median (line within box), 1st and 3rd quartiles (box), margined by largest and smallest data points (whiskers) (total 10 sessions, 5 left and 5 right infusions for each target). Local DCZ infusion into unilateral ventral putamen had no significant effects on performance (Brunner-Munzel test, BM-value = 0.44, p = 0.67). (**E,F**) Behavioral effects of local DCZ infusion into unilateral ventral putamen on delayed response task. Correct performance rate (mean ± sem, total 10 sessions, 5 left and 5 right infusions for each target) following DCZ (ipsilateral, magenta; contralateral, purple) and vehicle infusions (cyan). Local DCZ infusion into unilateral ventral putamen had no significant effect on performance in the two trial types (two-way ANOVA, main effect of treatment, ipsilateral; F_(1,24)_ = 0.23, p = 0.64, contralateral; F_(1,24)_ = 0.04, p = 0.84). Scale bars on MR images: 5 mm.

